# Heart Rate Variability During Mindful Breathing Meditation

**DOI:** 10.1101/2022.08.06.503051

**Authors:** Aravind Natarajan

## Abstract

In this article, we discuss Heart Rate Variability (HRV) measured during mindful breathing meditation. We provide a pedagogical computation of 2 commonly used HRV metrics, i.e. the root mean square of successive differences (RMSSD) and the standard deviation of RR intervals (SDRR), in terms of Fourier components. It is shown that the RMSSD preferentially weights higher frequency Fourier modes, making it unsuitable as a biosignal for mindful breathing meditation which encourages slow breathing. We propose a new metric called the autonomic balance index (ABI) which uses Respiratory Sinus Arrhythmia to quantify the fraction of HRV contributed by the parasympathetic nervous system. We apply this metric to HRV data collected during two different meditation techniques, and show that the ABI is significantly elevated during mindful breathing, making it a good signal for biofeedback during meditation sessions.

## I. INTRODUCTION

Mindfulness has shown promise as a non-pharmaceutical intervention in the management of stress, as well as a variety of other conditions^1–5^. Meditation and mindfulness practices have the ability to support individuals, especially during difficult times^6,7^. Mindful breathing exercises have shown promise in helping to reduce reactivity to repetitive thoughts^8^.

In a metastudy of 7 controlled and randomized controlled studies which were aggregated, mindfulness based stress reduction (MBSR) was shown to have a significant positive non-specific effect compared to the absence of any treatment when comparing Cohen’s *d* measures of stress^9^. A study involving 75 participants who engaged in an 8 week course on MBSR showed a significant (Cohen *d* = 1.04) decrease in stress as measured by the 10-item Perceived Stress Scale^10^. A study involving 53 participants who attended a 10-day Vipassana meditation retreat showed reductions in overall distress 3 months following the retreat, encompassing a spectrum of psychological symptoms^11^. Mindfulness based treatments are also pursued in the management of chronic pain^12–14^ and insomnia^15–19^.

Commercially available wearable devices are increasingly popular in the United States, and many wearable devices offer mindfulness training^20–25^. A notable technique for training in mindfulness meditation is in the use of biofeedback signals. Biofeedback can help individuals gain awareness of physiological processes occurring within the body, and also to consciously control those processes^26^. A promising biosignal is the heart rate variability (HRV) which refers to the beat-to-beat variability in heart rate. A high HRV usually indicates good health and an increased ability to adapt to stressful situations. HRV biofeedback has been applied to the management of stress^27^, depression^28,29^, and asthma^30^. In a meta-analytic review of HRV biofeedback, it was shown that HRV biofeedback produces improvement in a variety of physical and emotional conditions^31^.

HRV is one of the best non-invasive probes of the autonomic nervous system (ANS)^32^. The ANS consists of 2 main branches: the sympathetic branch which predominates during exercise, and stressful “fight or flight” reactions, and the parasympathetic branch which predominates during quiet, resting conditions^33^. The tenth cranial nerve called the vagus nerve is the main contributor of the parasympathetic nervous system (PNS) and the provides the main parasympathetic supply to the heart^34,35^. A valuable metric of vagal or parasympathetic activity is Respiratory Sinus Arrhythmia (RSA)^36–43^ which is the rhythmic modulation of the heart rate in response to respiration. The heart rate increases during inhalation, and decreases during exhalation, and this phenomenon has been associated with the efficiency of pulmonary gas exchange^38,44,45^. PNS activity may be utilized to quantify stress by defining stress as a disruption of homeostasis with low PNS activity^42^. A state characterized by the absence of stress would therefore be one with high PNS activity^42^. The PNS activity can be quantified by measuring the RSA which manifests as excess power in the HRV power spectrum, at the respiratory frequency.

The connection between HRV and meditative states of mind has been well established in the scientific literature. Murata et al.^46^ collected EEG data and HRV data during Zen meditation, and analyzed the data in association with trait anxiety. It was found that slow alpha wave inter-hemispheric EEG coherence in the frontal lobe increased during meditation, reflecting non-task related cognitive processes such as attention. Among HRV measures, this was accompanied by an increase in the relative HF power and decrease in LF/HF, reflecting an increased parasympathetic response (the respiratory rate was fixed to 15 per minute). Wu and Lo^47^ reported HRV changes among two groups: the first group consisting of 10 experienced Zen practitioners, while the other group consisting of non-meditators. They found that when the ANS was under parasympathetic predominance, the heart rate can be purely modulated by respiration (their respiratory rate was about 15 per minute). Nesvold et al.^48^ studied HRV changes during non-directive meditation, and found an increase in both LF and HF components (the respiration rate was unchanged, they interpret the change in HRV as entirely due to meditation, not changes in respiration). They also found no change in mean heart rate during meditation. Cysarz and Büssing^49^ investigated the impact of 4 exercises: spontaneous breathing, mental task, seated Zen, and walking meditation, on HRV. Seated Zen and walking meditation both resulted in a high degree of synchronization between respiration and heart rate, while spontaneous breathing and the mental task showed no such synchronization. The two kinds of meditation were characterized by increased LF (due to a much slower breathing rate) and in-phase RSA. Lo et al.^50^ studied the effect of Zen meditation on subjects undergoing a drug rehabilitation program, showing significant improvement in HRV (especially RSA), but no change in HR.

A popular HRV metric computed by many commercial wearable devices, and which is often regarded as a measure of the PNS is the Root Mean Squared value of the Successive Differences (RMSSD) of the interbeat intervals (henceforth “*RR* intervals”)^51^. This is indeed the case when the HRV is measured during sleep, when the respiratory rate is typically^52^ in the range 11.8 min^−1^ 19.2 min^−1^ and the RSA appears as excess power in the high frequency band (9 min^−1^ 24 min^−1^) of the HRV power spectrum. This is true because the RMSSD is a *biased* estimator of HRV, i.e. it preferentially weights high frequency components and is therefore, sensitive to RSA provided the respiratory rate is within the high frequency band. The RMSSD is not as informative about parasympathetic activity during slow paced breathing when the respiratory rate can be as low as 6 min^−1^ or even lower, and when the RSA falls within the low frequency band (2.4 min^−1^ 9 min^−1^).

In this article, we will provide a pedagogical calculation to demonstrate that the RMSSD should only be considered when the HRV is dominated by Fourier modes in the high frequency band, e.g. during sleep. When compared to an unbiased metric such as the Standard Deviation of the RR intervals (SDRR), we will see that the RMSSD greatly underestimates the HRV when the respiratory rate is low, i.e. during slow, paced breathing favored during mindful breathing meditation. The SDRR is however, a measure of the total ANS, and not the PNS. We will therefore consider another metric based on RSA called the *autonomic balance index* (ABI) which is the ratio of HRV due to respiration to the total HRV. When RSA is the dominant source of HRV, the *RR* interval time series resembles a periodic sine wave due to a small number of dominant Fourier components, a condition known as coherence (see for example, McCraty et al.^53^). Our computation of the ABI is qualitatively similar to the coherence ratio computation described in McCraty et al.^53^ (but the ABI is bounded between 0 −1). We hypothesize that the ABI is proportional to the ratio PNS:ANS, and which can therefore be interpreted as a measure of the absence of stress. Proxies of autonomic balance have been considered in the literature, e.g. LF/HF or Poincare *S*_1_*/S*_2_, however, these measures do not work during slow, mindful breathing. During slow breathing, the RSA falls within the LF band, and HF power does not capture PNS activity, rendering the LF:HF ratio unsuitable. The Poincare *S*_1_ and *S*_2_ parameters are linearly related to RMSSD and SDRR and hence, they too cannot be used during slow breathing. We will provide a simple algorithm to compute the ABI. We will then apply this computation to a publicly available dataset of HRV measured during meditation, and show that ABI is largest during mindful breathing.

## II. METHODS

### A. Data

The data used for this analysis have been described by Peng et al.^54^ and may be downloaded^55^ from the Physionet database^56^. Two specific meditative techniques were investigated by Peng et al.^54^: (i) Chinese Chi (Qigong) meditation and (ii) Kundalini Yoga meditation. There were 8 Chi meditators (5 female and 3 male, age range 26−35, mean 29 yr, with 1−3 months of prior practice) and 4 Kundalini Yoga meditators (2 female and 2 male, age range 20−52, mean 33 yr, advanced meditators). Time series data of the instantaneous heart rate have been provided from which we computed the *RR* interval time series data. Data were collected during meditation and also during the period prior to meditation, which serves as a control. Also included were three additional non-meditating cohorts to serve as an additional control: (iii) 14 healthy subjects (9 female, 5 male, age range 20−35, mean 25 yr, supine) following metronomic breathing at 15 min^−1^, (iv) 11 healthy subjects (8 female, 3 male, age range 20−35, mean 29 yr) during sleep, and (v) 9 elite triathlon athletes (3 female, 6 male, age range 21−55, mean 39 yr) during sleep. This is summarized in Table I, ‘Duration’ refers to the average duration (minutes) of data per volunteer, and *N*_5_ is the total number of 5-minute segments we used in the analysis.

**TABLE I.** 7 cohorts of volunteers: Chi (med) and Yoga (med) are the two meditation cohorts, while Chi (rest) and Yoga (rest) are controls prior to meditation. Also considered are the metronomic breathing, normal healthy adults during sleep, and elite athletes during sleep. ‘Duration’ refers to the average time per volunteer (minutes), and *N*_5_ is the total number of 5-minute segments we used in the analysis.

| Cohort | Description | Volunteers | Duration (min.) | $N_5$ |
| --- | --- | --- | --- | --- |
| Chi (med) | Chi meditation | 8 | 57.1 | 67 |
| Chi (rest) | Prior to meditation | 8 | 58.3 | 54 |
| Yoga (med) | Kundalini Yoga meditation | 4 | 11.1 | 8 |
| Yoga (rest) | Prior to meditation | 4 | 10.1 | 6 |
| Metronomic | Metronomic breathing | 14 | 10.0 | 25 |
| Sleep - Normal | Sleep - healthy individuals | 11 | 352.6 | 535 |
| Sleep - Ironman | Sleep - elite athletes | 9 | 85.9 | 93 |

### B. Quantifying Respiratory Sinus Arrhythmia

Let us now investigate a technique for quantifying the autonomic balance through RSA. There have been multiple methods suggested for quantifying RSA (see for e.g. Ref.^57^). In this article, we consider a different technique similar to well known HRV metrics, and one which is suitable for use during slow, paced breathing, especially when the condition of coherence is attained. McCraty et al.^53^ quantify the condition of coherence through the coherence ratio. Here we will consider a similar approach: We first compute the standard deviation of the RR intervals due to RSA alone, called the SDRSA, considered to be a probe of the PNS. We then compute the ratio (SDRSA / SDRR) which is bounded between 0 − 1, and which we refer to as the Autonomic Balance Index (ABI).

A required step in computing the SDRSA is the measurement of the respiratory rate. A possible complication here is that the respiratory rate is variable when the subject is awake. We therefore consider segments that are short enough that we may make the assumption that the respiratory rate is approximately constant within that segment. It is also important that the chosen segment is not too short because (i) a very short time window will admit only a small number of realizations of each Fourier mode, increasing the shot noise error, and (ii) the resolution in the spectral domain is inversely proportional to the size of the window in the time domain. We choose a segment size of 2 minutes and smooth the signal with a Hann window. Estimation of SDRSA and ABI follow the algorithm:

1. Define 2 frequencies *f*_1_ and *f*_2_ that may be considered the lower and upper bounds for the respiratory rate within each 2 minute segment.
2. Compute the power spectral density (PSD) normalized so that ∫ *df P* (*f*) = SDRR^2^. The PSD is interpolated using a cubic spline. Let *f*_0_ be the frequency that corresponds to the peak of the PSD, and let *A*_0_ be the peak value.
3. Fit a Gaussian 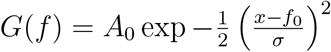 to the peak of the PSD described by a mean value (*f*_0_), a standard deviation (*σ*), and amplitude *A*_0_.
4. Construct the residual *R*(*f*) = *PSD*(*f*) −*G*(*f*). From the residual, identify the largest peak amplitude *A*_1_ in the range *f*_1_ *< f < f*_2_. Compute the ratio *P* = *A*_0_*/A*_1_ which represents the prominence of the main peak *A*_0_. If *P* is greater than a preset limit *P*_min_, it validates our assumption that the PSD is dominated by a single respiratory frequency. If *P < P*_min_, no values are returned and the data are discarded.
5. If *P > P*_min_, we compute the following two quantities: (i) The variance due to respiration = SDRSA^2^, estimated by the area under the gaussian 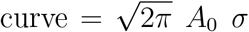. (ii) The normalized quantity ABI = [SDRSA*/*SDRR]. The algorithm returns the estimated respiratory rate *f*_0_ and ABI.

We apply the algorithm above during meditation, metronomic breathing, and sleep. For the data measured during rest prior to meditation (Chi (rest) and Yoga (rest)) however, we include an additional step at the beginning because the respiratory rate is highly variable, and the algorithm works best when the respiratory range is fairly small. We compute the frequency *f*_0_ corresponding to the peak of the PSD and set *f*_1_ = MAX(12 min^−1^, *f*_0_ − 3 min^−1^) and *f*_2_ = MIN(22 min^−1^, *f*_0_ + 3 min^−1^). For the meditation cohorts (Chi (med) and Yoga (med)), we set *f*_1_ = 3 min^−1^ and *f*_2_ = 10 min^−1^. For the metronomic, normal, and ironman cohorts, we set *f*_1_ = 10 min^−1^ and *f*_2_ = 20 min^−1^. In all cases, we set *P*_min_ = 2.

The data are initially divided into non-overlapping 5 minute segments. Each 5 minute segment is then divided into a number of 2 minute segments with an overlap of 10 seconds. To ensure sufficient data for analysis, we estimate the coverage in each 2 minute segment as the number of observed heart beats / expected number of heart beats, and impose the condition that the coverage *>* 0.7. Provided the coverage condition is met, *f*_0_ and ABI are estimated from each such 2 minute segment (starting from 2:00, in increments of 10 seconds, until the 5:00 minute mark). A total of 19 such estimates can be made from a 5 minute segment of data (from 2:00 to 5:00 in increments of 10s, including both endpoints). The median value of these different estimates is then calculated for *f*_0_ and ABI provided there is a minimum of 9 estimates. If there are fewer than 9 estimates (for example, due to missing data), we do not store any results for that 5 minute segment.

## III. RESULTS

### A. An analytic approximation for the RMSSD and SDRR

As mentioned in the Introduction, the RMSSD is influenced by the respiratory rate and is therefore hard to interpret. Here, we provide a pedagogical approximation of the RMSSD and SDRR from first principles, using example data measured during slow breathing.

Let *RR*_*i*_ represent the *i*^th^ value of the *RR* interval time series sampled at time intervals *t*_*i*_ = [*t*_1_, *t*_2_, *t*_3_, …], where *RR*_*i*_ = *t*_*i*_ − *t*_*i*−1_. The *RR* time series contains a constant term ⟨*RR*⟩ and a fluctuating term 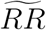

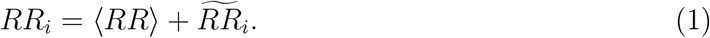

The standard deviation of the RR intervals (SDRR) is computed as:

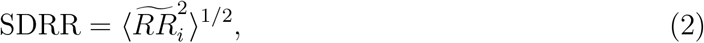

where the angle brackets represent the mean value. The fluctuating component is expected to be much smaller than the mean, i.e. SDRR ≪ ⟨*RR*⟩. The differences between successive *RR* intervals Δ*RR*_*i*_ may be computed as:

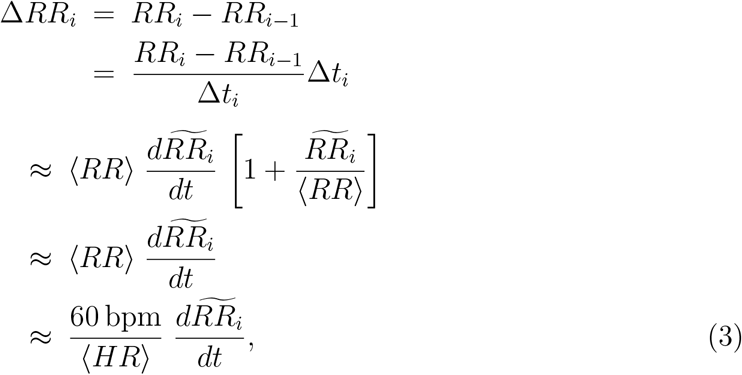

where we used Δ*t*_*i*_ = *t*_*i*_ − *t*_*i*−1_ = *RR*_*i*_, and we ignored term of quadratic order in 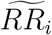. ⟨*HR*⟩ is the mean heart rate, and “bpm” stands for beats per minute. The RMSSD is the root mean square of the successive differences Δ*RR*_*i*_, i.e.

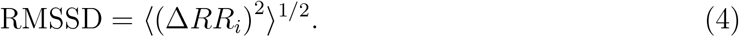

Let us interpolate and re-sample the 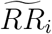 sequence at a sampling frequency *N/T*_0_ to obtain an 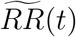 field with *N* samples (and where *T*_0_ is the length of the signal under consideration, we will assume that the end points are identified to mimic periodicity). We may expand this in a Fourier series:

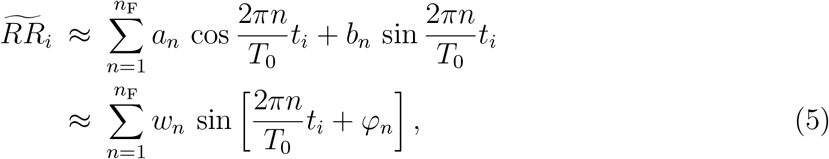

where *n*_F_ ≤ *n*_max_ is the total number of Fourier modes to be included in the approximation, and by Nyquist’s theorem, *n*_max_ = *N/*2. 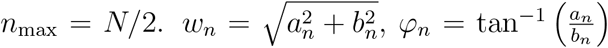, and *t*_*i*_ are the time intervals at which the *RR* intervals are calculated. We have assumed *b*_*n*_ ≠ 0 and 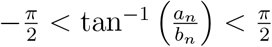. The coefficients *a*_*n*_ and *b*_*n*_ may be computed as:

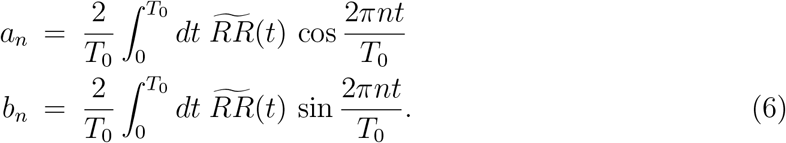

The frequencies of the Fourier modes are 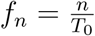 for *n* = 1, 2, 3, … *n*_F_. Let SDRR(*n*_F_) and RMSSD(*n*_F_) be the values of SDRR and RMSSD estimated using *n*_F_ ≤ *n*_max_ Fourier modes, while SDRR and RMSSD are the exact values. Eq. 5 can be used to estimate SDRR(*n*_F_) from Eq. 2. From Eq. 5 and Eq. 3, we find

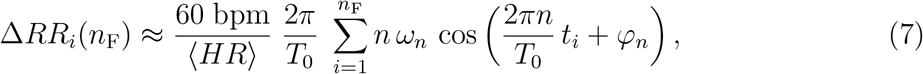

from which we can estimate the RMSSD(*n*_F_) using Eq. 4.

Fig. 1(a) shows a 5 minute sample of RR intervals from a participant practicing Chi meditation, exhibiting high HRV and prominent RSA. Plot (b) shows the power spectral density along with the best fit gaussian curve. We note that most of the power comes from a narrow band of frequencies centered around a respiratory frequency of 3.2 min^−1^. Plot (c) shows the ratios [RMSSD (*n*_F_) / RMSSD] and [SDRR (*n*_F_) / SDRR], where the values of RMSSD (*n*_F_) and SDRR (*n*_F_) are calculated from the approximate formula we derived, and include up to *n*_F_ Fourier modes. RMSSD and SDRR are the exact values. Both RMSSD (*n*_F_) and SDRR (*n*_F_) approach their true values as *n*_F_ → *n*_max_, but the RMSSD (*n*_F_) computation made some simplifying assumptions, and is hence not as accurate as the SDRR (*n*_F_) computation. RMSSD (*n*_F_) also converges to the true value with a far larger number of Fourier modes than SDRR (*n*_F_) since it preferentially weights high frequency modes. SDRR (*n*_F_) requires only 18 Fourier modes (corresponding to a peak frequency of 88/5 = 3.6 min^−1^) to reach 90% of the true value. This makes intuitive sense since the peak of the PSD was found to be at 3.2 min^−1^ and there is very little power at higher frequencies. RMSSD (*n*_F_) on the other hand, requires 88 Fourier modes (corresponding to a peak frequency of 88/5 = 17.6 min^−1^) to reach 90% of the true value. Frequencies above 3.6 min^−1^ contribute ≲ 10% to the SDRR, while contributing ≈ 35% to the RMSSD. This discussion highlights a major flaw in using the RMSSD to quantify HRV for slow breathing:

**FIG. 1.**
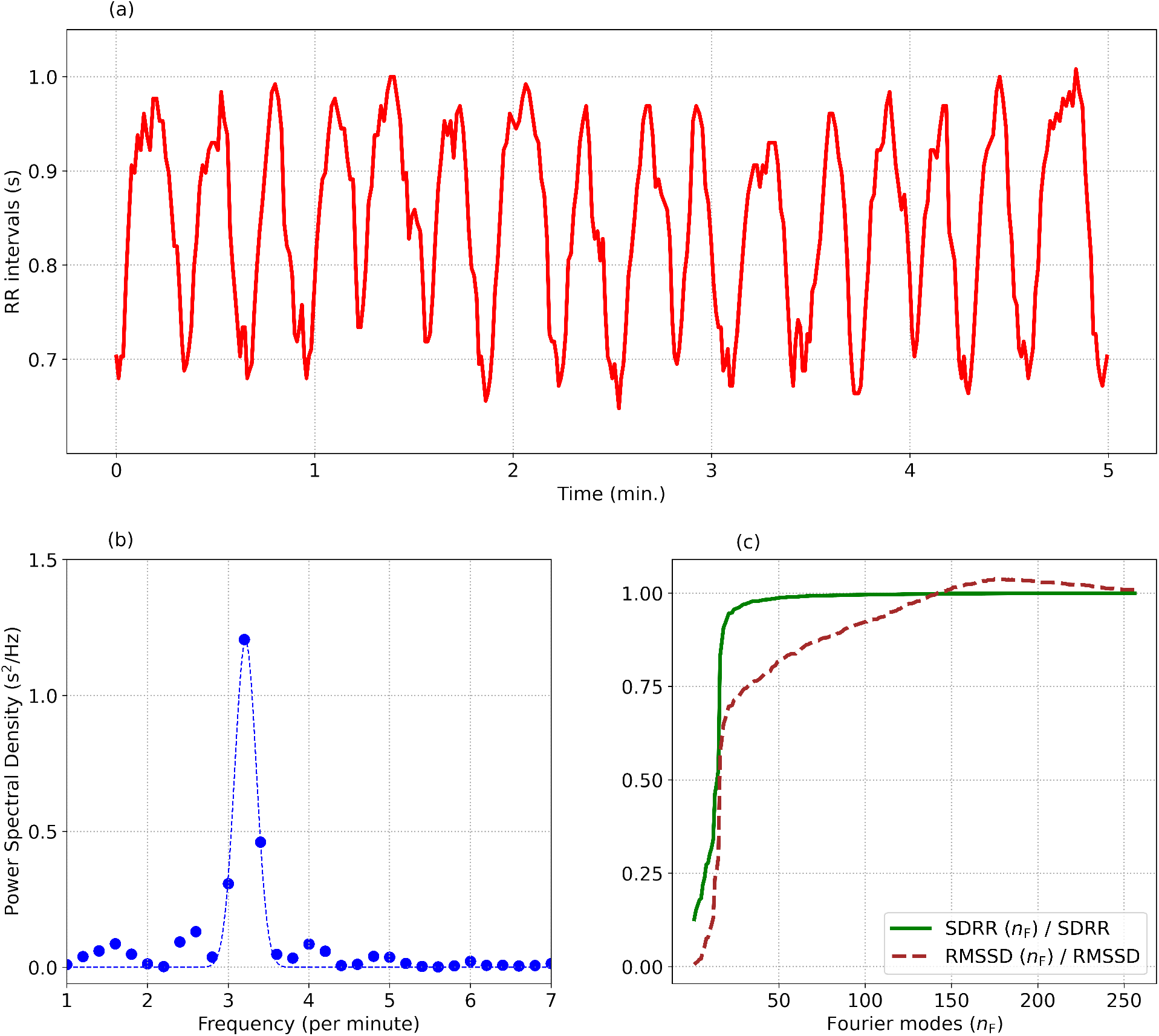
(a) describes a 5 minute segment of the *RR* interval time series, from a subject practicing Chi meditation. (b) shows the power spectral density showing strong respiratory sinus arrhythmia at a respiratory frequency of 3.2 min^−1^. The ratios [RMSSD(*n*_F_)/RMSSD] and [SDRR(*n*_F_)/SDRR] computed from the first *n*_F_ Fourier modes are displayed in (c). Many more Fourier modes need to be included for the computation of RMSSD (*n*_F_) compared to SDRR (*n*_F_) since RMSSD preferentially weights the higher frequency modes.

A small amount of power at high frequencies is preferentially weighted by the RMSSD even though the high frequency Fourier modes in this case are not associated with respiration. The SDRR being an unbiased HRV estimator does not weight Fourier modes differently by frequency. The lack of high frequency content also results in the RMSSD being much lower than the SDRR. For this example, we find RMSSD = 36.8 ms, while the SDRR = 102.6 ms. Fig. 1 demonstrates a sample wherein most of the variance is contributed by a single Fourier mode (or a narrow range of modes). Let us consider the special case when all the power comes from a single frequency mode. The *RR* time series may then be simplified as:

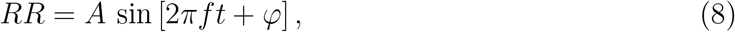

where *f* is the respiratory rate and *A* is the amplitude of oscillations. The SDRR is then simply 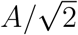. We can approximate Δ*RR* as:

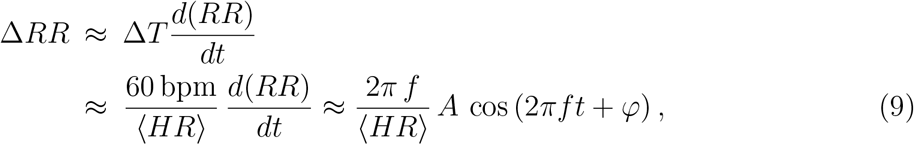

where *f* is measured in min^−1^ and ⟨*HR*⟩ is measured in beats per minute. Taking the root mean square of Eq. 9, we get:

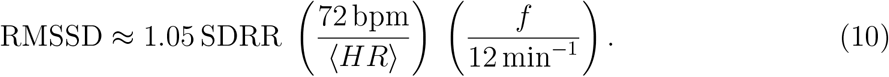

It is clear from Eq. 10 that the RMSSD increases with respiratory rate. This metric is therefore best employed in situations when the respiratory rate is high (i.e. *>* 12 min^−1^) and relatively constant, i.e. during sleep. In the example shown in Fig. 1, we find the respiratory rate = 3.2 min^−1^, and the mean heart rate is 72 beats per minute. From our approximate analysis (Eq. 10), we expect a ratio RMSSD:SDRR = 0.28, while the true ratio = 0.36.

### B. Autonomic balance during meditation

We have seen in the previous subsection that the RMSSD is unsuitable during periods of slow, paced breathing favored by mindful breathing meditation. The SDRR, although an unbiased metric, is also not ideal for biofeedback during mindful breathing meditation since it captures the total variance, i.e. it is a measure of the ANS and not the PNS. In this subsection, we apply the algorithm for computing ABI described in the Methods section, to the data.

Fig. 2 shows the mean subtracted RR interval time series data for 2 situations: (a) describes a 2 minute resting period prior to meditation, while (c) shows the data during meditation (we used an sample from the Kundalini Yoga cohort for this figure). (b) and (d) show the power spectral density plots for the 2 situations respectively. Contrasting the 2 scenarios, we note the following: (i) The respiratory rate is much lower during the meditation phase (4 min^−1^) compared to the resting phase (18.5 min^−1^), (ii) Most of the power is contained within the respiratory band during the meditation phase (ABI = 0.86). In the case of the resting period prior to meditation, there is considerable power at frequencies not associated with respiration (ABI = 0.57, some of this power is likely due to Mayer Wave Sinus Arrhythmia^58,59^), (iii) The amplitude of oscillations is much larger during meditation (SDRR = 55.7 ms) compared to the resting phase (SDRR = 26.4 ms).

**FIG. 2.**
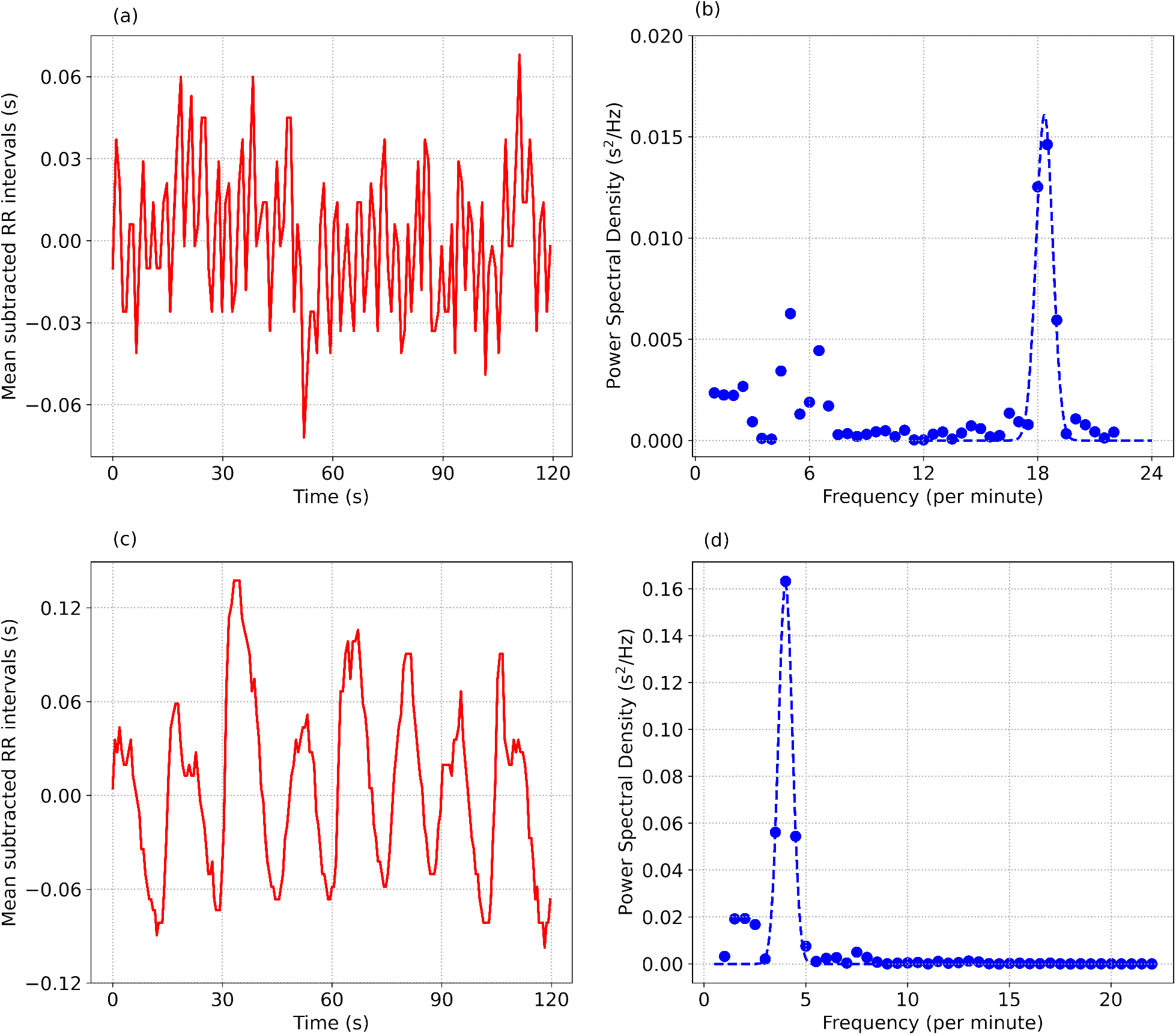
(a) and (b) show the mean subtracted RR interval time series data and the power spectral density, from a 2 minute segment of data collected prior to meditation. (c) and (d) show the same quantities for data collected during Kundalini Yoga meditation.

Fig. 3 shows various metrics evaluated for the 7 different cohorts: (i) Chi (med), i.e. during Chi meditation, (ii) Chi (rest), i.e. prior to meditation, (iii) Yoga (med) during Kundalini Yoga meditation, (iv) Yoga (rest) prior to meditation, (v) Metronomic breathing, (vi) Normal, i.e. healthy individuals during sleep, and (vii) Ironman triathletes during sleep. The Chi cohort (meditation or rest) is shown in magenta, the Yoga cohort (meditation and rest) in green, metronomic breathing in red, normal in brown, and ironman in blue. Subplot (a) shows the ABI for the 7 cohorts. The highest scores are obtained for the 2 meditation cohorts, followed by metronomic breathing, sleep, and finally the rest cohort (awake, but not meditating). Subplot (b) shows the RMSSD. Not surprisingly the mediation cohorts perform poorly in the RMSSD comparison due to the dependence of RMSSD on respiratory rate. Subplot (c) shows the average heart rate. There is very little difference in heart rate during Chi meditation. For the Kundalini Yoga cohort however, the heart rate *increases* during meditation. The heart rate is lowest during sleep, especially for the elite athletes. We note the heart rate is not expected to decrease significantly during mindful breathing^32^, and is therefore not a good metric to use as biofeedback. Subplot (d) shows the respiratory rate. We see that the 2 meditation techniques we discuss here encourage very slow breathing. As expected the metronomic breathing cohort shows very little variability.

**FIG. 3.**
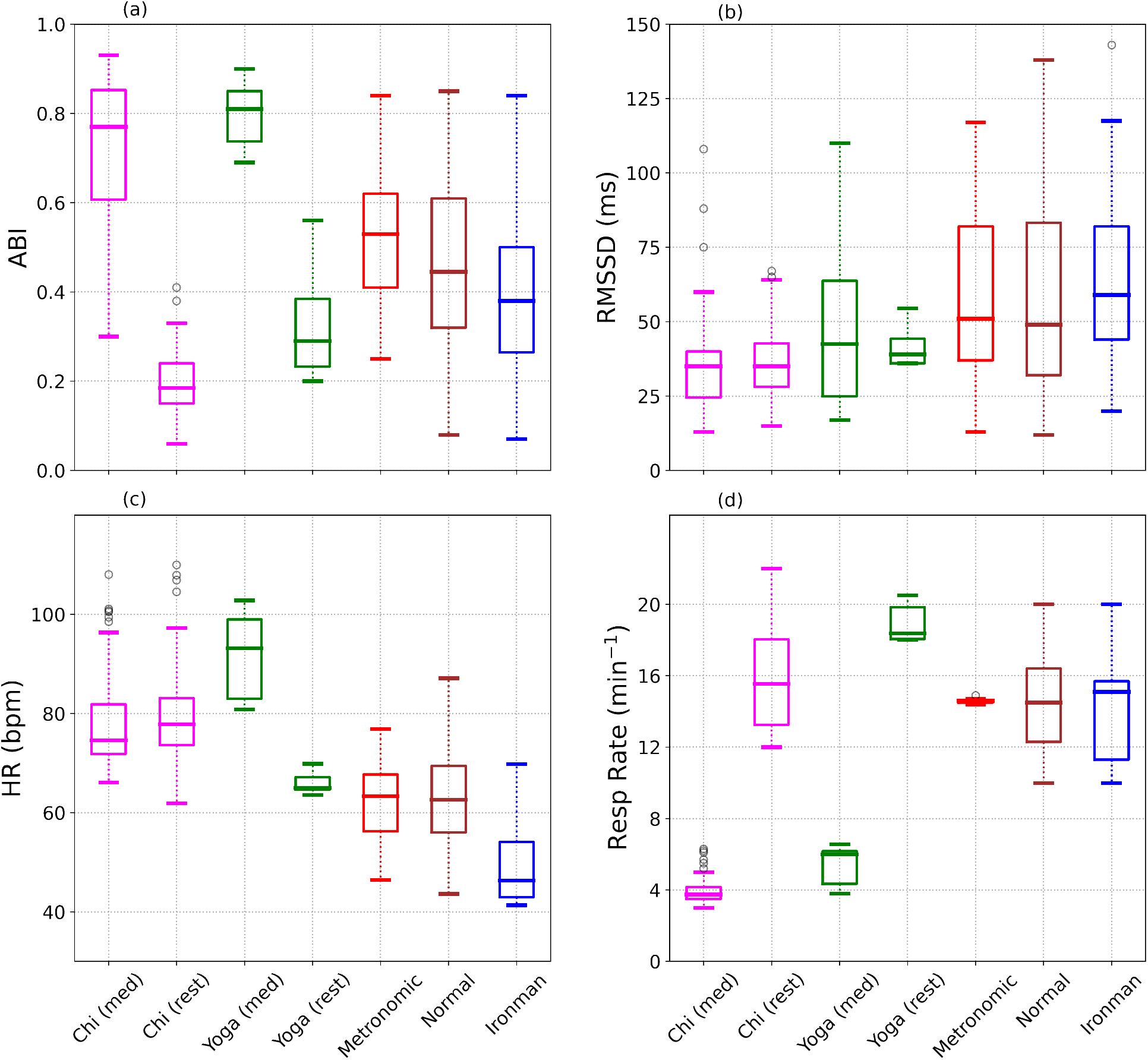
A comparison of the autonomic balance index (ABI), RMSSD, Heart Rate, and Respiratory Rate for the 7 different cohorts. ABI shows a very significant difference between the meditation and rest data, while such a difference is not seen in the RMSSD. The respiratory rate is also substantially decreased during meditation compared to the rest phase.

Table II shows the mean (standard deviation) computed from the 5-minute medians, for the autonomic balance index ABI, respiratory rate, RMSSD, SDRR, and heart rate (HR) for the seven different cohorts. One can compare a pair of cohorts by means of the Cohen *d* effect size^60,61^:

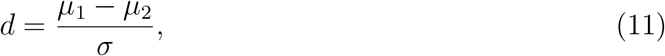

where *μ*_1_ and *μ*_2_ are the means of the 2 cohorts, and *σ* is the pooled standard deviation given by:

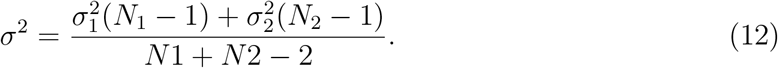

Compared to the Chi (rest) cohort, the Chi (med) cohort achieved a higher ABI with an effect size *d* = 3.90. The Yoga (med) cohort showed a higher ABI compared to the Yoga (rest) cohort, quantified by the effect size *d* = 4.82. The metronomic breathing cohort showed a slightly higher ABI compared to the normal cohort (*d* = 0.38) and the ironman cohort (*d* = 0.67). For the RMSSD metric, the Chi (med) and Chi (rest) cohorts were found to be similar (*d* = −0.06), while the Yoga(med) showed a slightly higher RMSSD than Yoga (rest) (*d* = 0.37). The difference between the meditation and rest cohorts are more clear in the SDRR. The Chi (med) cohort was characterized by a higher SDRR compared to the Chi (rest) cohort (*d* = 0.52), while the Yoga (med) cohort had a higher SDRR compared to the Yoga (rest) cohort (*d* = 1.11). This is expected from our earlier discussion on RMSSD being lowered in cases of low respiratory rate, while the SDRR is unaffected by the respiratory rate. Chi (med) showed a slightly decreased heart rate compared to Chi (rest) (*d* = −0.19), while Yoga (med) interestingly showed a significant *increase* in heart rate compared to Yoga (rest) (*d* = 3.9). The cohort with the lowest heart rate were the elite athletes: the triathlon cohort was found to have a lower heart rate compared to the metronomic breathing cohort (*d* = −1.68) as well as the normal, healthy cohort (*d* = −1.44). When comparing the respiratory rate, the 2 meditation cohorts have the lowest rates: The respiratory rate for the Chi (med) cohort was much lower than the Chi (rest) cohort (*d* = −6.07). The Yoga (med) cohort similarly showed a much lower respiratory rate compared to the Yoga (rest) cohort (*d* = −12.27).

**TABLE II.** Mean (standard deviation) computed for the 7 cohorts, for different metrics.

| Activity | ABI | Resp. Rate<br>(min<sup>-1</sup>) | RMSSD<br>(ms) | SDRR<br>(ms) | HR<br>(bpm) |
| --- | --- | --- | --- | --- | --- |
| Chi (med) | 0.72 (0.17) | 3.9 (0.8) | 35.5 (16.3) | 66.9 (22.5) | 77.8 (9.7) |
| Chi (rest) | 0.19 (0.07) | 15.8 (2.8) | 36.4 (12.8) | 56.8 (14.7) | 79.7 (10.1) |
| Yoga (med) | 0.80 (0.07) | 5.4 (1.1) | 50.4 (31) | 86.6 (32.3) | 91.9 (8.5) |
| Yoga (rest) | 0.33 (0.12) | 18.9 (1.1) | 41.6 (6.7) | 56.8 (16) | 66 (2.2) |
| Metron | 0.52 (0.15) | 14.6 (0.1) | 59.1 (30.2) | 58.1 (20.6) | 61.7 (8.1) |
| Normal | 0.45 (0.18) | 14.6 (2.7) | 58.1 (32.2) | 67.9 (39.8) | 62.1 (9.1) |
| Ironman | 0.40 (0.18) | 14.1 (2.4) | 63.5 (25.4) | 78.8 (46.5) | 49.4 (7.1) |

## IV. DISCUSSION

In this article, we discussed heart rate variability measured during mindful breathing meditation. We first considered the RMSSD and SDRR, two popular HRV metrics used by commercial wearable devices to quantify HRV. We derived an approximate but pedagogical, analytic expression for SDRR(*n*_F_) and RMSSD(*n*_F_) using Fourier decomposition, and including the first *n*_F_ number of Fourier modes. This pedagogical exposition made it clear that the RMSSD is a *biased estimator* of the HRV in that it preferentially weights higher frequency Fourier components, with the result that a small amount of power at high frequencies can contribute a disproportionately large influence on the RMSSD. Such an effect is not seen in the SDRR which weights all Fourier modes equally. RMSSD is thus, not a suitable metric to quantify HRV during slow, mindful breathing.

We have suggested a metric that quantifies the fraction of HRV contributed by the RSA as a HRV metric that is ideally suited to serve as a biofeedback signal during mindful breathing meditation. The ABI metric was motivated by the spectral properties of HRV during mindful breathing, and is qualitatively similar to the coherence ratio computation described in McCraty et al.^53^. During mindful breathing, most of the power falls within the respiratory band of frequencies, with very little power at lower frequencies (note that the respiratory frequency itself may be as low as ∼ 3 min^−1^) indicating that most of the HRV is due to respiratory sinus arrhythmia. Unlike other HRV measures, ABI is less influenced by age, gender, physical fitness etc as it is a ratio of 2 HRV measures. Instead, it is most influenced by practices that result in PNS dominance, e.g. meditation. We described a simple algorithm to compute ABI from the power spectral density of *RR* fluctuations.

We then applied the ABI to heart rate time series data collected during meditation, and described in Peng et al.^54^. The authors^54^ found extremely prominent HRV fluctuations during 2 specific, traditional meditation techniques: Chinese Chi and Kundalini Yoga (here denoted as Chi (med) and Yoga (med) respectively). The data also included a period of rest prior to meditation (here denoted as Chi (rest) and Yoga (rest) respectively). As additional controls, the data also included metronomic breathing (“metronomic”), healthy adults during sleep (“normal”), and elite athletes during sleep (“ironman”). The values of ABI and RMSSD for the 7 different cohorts are shown in Fig. 3(a) and Fig. 3(b) and demonstrate that ABI is a more sensitive metric than the RMSSD. The mean and standard deviation of ABI, Respiratory Rate, RMSSD, SDRR, and heart rate are tabulated for the 7 cohorts in Table II. ABI and the respiratory rate are significantly different in the mediation cohorts compared to the others. We therefore recommend ABI and respiratory rate as potential biosignals to assist with meditation practice.

There are several limitations to this work. The ABI algorithm interprets power at frequencies outside the chosen range (*f*_1_, *f*_2_) of possible respiratory rates as due to stress, i.e. sympathetic nervous system activity creates low frequency power. However not all power at low frequencies is due to stress, e.g. the Mayer Wave Sinus Arrhythmia (MWSA) at *f* ≈ 6 per minute is caused by blood pressure oscillations^58,59^, and a pronounced MWSA can cause an artificially low ABI when the RSA occurs at much higher frequencies. Another limitation in computing the ABI is that it assumes a constant respiratory rate within a 2 minute window. While this is naturally satisfied during mindful breathing or during sleep, that is less likely to be the case when subjects are awake. We have also assumed that the inhalation and exhalation times are the same. ABI also relies on the presence of respiratory sinus arrhythmia. We were able to compute ABI in the period prior to meditation, but we expect it would be harder to do so at a random time of day, and especially during times of stress when the respiratory sinus arrhythmia would be subdominant. RSA is also decreased in older individuals and it would therefore be harder to compute ABI for older subjects. The algorithm is unlikely to work in individuals who have heart arrhythmias. ABI would likely not be computed during normal activities such as working, eating, watching television, etc, and other HRV metrics would be preferable during these activities. Wearable devices also have difficulty measuring *RR* interval data when subjects are moving.

## VI. DATA SHARING

All data may be downloaded from https://physionet.org/content/meditation/1.0.0/.

## VII. ACKNOWLEDGMENTS

I acknowledge funding from Fitbit Research, Google LLC. It is a pleasure to thank members of the Fitbit Research team for many helpful discussions.

